# Blood transcriptomics analysis offers insights into variant-specific immune response to SARS-CoV-2

**DOI:** 10.1101/2023.11.03.564190

**Authors:** Markus Hoffmann, Lina-Liv Willruth, Alexander Dietrich, Hye Kyung Lee, Ludwig Knabl, Nico Trummer, Jan Baumbach, Priscilla A. Furth, Lothar Hennighausen, Markus List

## Abstract

Bulk RNA sequencing (RNA-seq) of blood is typically used for gene expression analysis in biomedical research but is still rarely used in clinical practice. In this study, we argue that RNA-seq should be considered a routine diagnostic tool, as it offers not only insights into aberrant gene expression and splicing but also delivers additional readouts on immune cell type composition as well as B-cell and T-cell receptor (BCR/TCR) repertoires. We demonstrate that RNA-seq offers vital insights into a patient’s immune status via integrative analysis of RNA-seq data from patients infected with various SARS-CoV-2 variants (in total 240 samples with up to 200 million reads sequencing depth). We compare the results of computational cell-type deconvolution methods (e.g., MCP-counter, xCell, EPIC, quanTIseq) to complete blood count data, the current gold standard in clinical practice. We observe varying levels of lymphocyte depletion and significant differences in neutrophil levels between SARS-CoV-2 variants. Additionally, we identify B and T cell receptor (BCR/TCR) sequences using the tools MiXCR and TRUST4 to show that - combined with sequence alignments and pBLAST - they could be used to classify a patient’s disease. Finally, we investigated the sequencing depth required for such analyses and concluded that 10 million reads per sample is sufficient. In conclusion, our study reveals that computational cell-type deconvolution and BCR/TCR methods using bulk RNA-seq analyses can supplement missing CBC data and offer insights into immune responses, disease severity, and pathogen-specific immunity, all achievable with a sequencing depth of 10 million reads per sample.

**Key Points:**

1. Computational deconvolution of transcriptomes can estimate immune cell abundances in SARS-CoV-2 patients, supplementing missing CBC data.
2. 10 million RNA sequencing reads per sample suffice for analyzing immune responses and disease severity, including BCR/TCR identification.

## Introduction

Peripheral blood is the tissue of choice in clinical diagnostics and biomedical research due to minimally invasive sample collection. As blood perfuses all organs, it provides insights into various diseases and medical conditions^1,2^. In general, we can investigate active pathways and organismal responses to stimuli (e.g., a viral infection) on a transcriptomic level^3,4^ by using well-established sequencing techniques such as bulk RNA sequencing (RNA-seq). A blood sample contains various cell types with different expression profiles. Complete blood counts (CBCs) are routinely assessed in the clinical setting and provide specific information regarding the proportions of cells present^5^. Within the white blood cell compartment, the percentages of neutrophils, lymphocytes, monocytes, eosinophils, and basophils provide insight into the type and response to infection and underlying disease and/or therapy^6–14^. However, CBCs are frequently unavailable in publicly accessible datasets, limiting insights into the status of the immune system.

Employing either bulk RNA-seq or single-cell RNA-seq (scRNA-seq, i.e., profiling gene expression at the individual cell level)^15^ can provide a detailed description of cellular composition, all based on the expression levels of genes. Performing scRNA-seq on each patient and cell type is not feasible due to the logistics and high costs of scRNA-seq^16^. To gain insights into the individual immune reactions to disease from RNA-seq data alone, it is essential to determine the composition of immune cells and gene expression in patient samples. CBCs - if available - provide a fundamental understanding of changes in the immune system; however, they do not specify more fine-grained segmentation into functional subgroups, which often drive disease progression. Hence, computational techniques such as MCP-counter^17^, xCell^18^, EPIC^19^, and quanTIseq^20^ (see Materials and Methods and Suppl. Materials 1 and 2 for differences, strengths, and weaknesses of each program) deconvolute bulk RNA-seq using signatures or gene sets of cell-type specific genes. They give a robust estimate of the abundance of various immune cell types within and across patient samples. Such insights into the status of an immune system are helpful for diagnostics, prognosis, and treatment selection with demonstrated potential in oncology^21^ and other diseases^10,12^. Here, we compare four different deconvolution approaches using bulk RNAseq to analyze changes in the white blood cell compartment over time in individuals infected with SARS-CoV-2.

In this study, we (1) compared computationally estimated immune cell abundances to CBC counts, the current gold standard. Moreover, we (2) investigate the immune cell abundances in patients infected with SARS-CoV-2 variants that differed in severity and tracked their progression over time, comparing them to a baseline model (i.e., seronegative samples taken from individuals that reportedly were never infected with SARS-CoV-2) to elucidate immune response differences and their progression over time to a healthy state. Additionally (3), we characterized the BCR and TCR profiles in infected patients. Finally (4), we compare how the performance of these methods is influenced by sequencing depth, i.e., how many reads have been sequenced for each sample.

## Methods

### Datasets

We utilized publicly available data from human buffy coat white blood cells from four distinct bulk RNA-seq experiments: GSE190680 (variants: Alpha, Alpha+EK (i.e., Alpha with an additional E484K mutation in the spike protein), Gamma)^22^, GSE162562 (seronegative)^23^, GSE201530 (variant: Omicron BA.1)^24^, and GSE205244 (variants: Omicron BA.1 and Omicron BA.2)^25^ (Suppl. Table 1a). Variants had samplings of days 0-5, 6-10, 11-15, 16-30, and >30 after hospitalization or onset of symptoms (Suppl. Tables 1a-b). All samples were processed by nf-core RNA-seq v. 3.8.1 using default parameters^26^. All 252 samples were controlled for quality by utilizing the reports of FastQC^27^ and MultiQC^28^, and only those (240 samples in total) with sufficient quality were included in the subsequent analyses (Suppl. Table 1b, Suppl. Table 2)^29,30^. Samples came from different studies but were processed in the same laboratory and with the same staff to avoid technical differences^31^.

### Immune deconvolution methodology

Cell-type deconvolution is a computational method applied to bulk RNA-seq data to estimate the abundance of cell types in a biological sample and is primarily used in the context of immune cells. In this study, we employ several tools bundled in the immunedeconv tool (using default settings established there), as it was previously shown that no single tool generally outperforms all others across all immune cell types^32^ (for marker genes See Suppl. Figs. 1a,b and https://doi.org/10.6084/m9.figshare.24442423.v1). Computational cell type deconvolution methods generally produce fractions or scores representing the abundance of specific cell types in the samples, which we use here for inter-sample comparisons between patients infected with different SARS-CoV-2 variants (see Suppl. Materials 1^17–20^).

### BCR/TCR repertoire methodology

BCR/TCR repertoire analysis refers to the study of the diverse collection of BCRs and TCRs present within an individual’s B and T cell repertoire (i.e., all unique antigen-specific receptors expressed on the surface of T cells and B cells), respectively. These receptors play a crucial role in the adaptive immune system by recognizing and binding to specific antigens derived from pathogens or abnormal cells, thus serving as biomarkers for past or current infections. Each BCR and TCR has a unique amino acid sequence, which contributes to the vast diversity and specificity of the immune response. We used two methods - MiXCR^33^ and TRUST4^34^ - to investigate bulk RNA-seq data by reconstructing B and T cell repertoires (Suppl. Materials 2^35–37^).

We used the Python package scirpy^38^ to analyze results from both methods. To extract only BCR/TCR sequences that differ from those found in a healthy population, we utilized the following steps: we computed a pairwise distance matrix for input sequences to identify sequences forming clonotypes and likely targeting similar antigens. Our objective was to identify sequences targeting SARS-CoV-2 antigens, enabling us to determine which BCR and TCR sequences respond to the virus. As sequences present in seronegative samples cannot target the SARS-CoV-2 virus, we can disregard them and focus on sequences exclusive to infected patients, further refining our search for the specific anti-SARS-CoV-2 receptor sequence. We use the ClustalW algorithm^39^ to perform multiple sequence alignment (see Suppl. Materials 3^40–43^). In the final steps, we employed the protein BLAST (pBLAST) tool^44,45^ to annotate clusters and sequences (e.g., those surpassing the cutoff and unlinked to sequences originating from healthy samples) to associate discovered sequences with specific viruses.

### Subsampling to a lower sequencing depth

RNA-seq always comes with a tradeoff between costs and information gain. Given the high sequencing depth (up to 150 million reads per sample) of the samples investigated here, we were interested in establishing a lower bound for obtaining robust results. To this end, we downsampled the samples to fifty and ten million reads using samtools^46^. Subsequently, we repeated both the immune deconvolution analysis and the BCR/TCR analyses. First, we generated TPMs expression matrices from the downsampled FASTQ files using salmon^47^ and revisited the immune deconvolution methods, then executed MiXCR and TRUST4 on the downsampled data and performed the subsequent analyses as described above.

## Results

In this study, we highlight the potential of RNA-seq data in clinical practice. Typically used for studying gene expression, RNA-seq data offers crucial insights into the status of the immune system via computational cell type deconvolution as well as the analysis of BCR and TCR sequences. While such advanced analysis techniques are increasingly widespread in oncology, we focus here on demonstrating their applicability in infectious diseases by example of SARS-CoV-2 infection. We re-analyzed data from 240 SARS-CoV-2 patients over time from the initial hospitalization through recovery^22,24,25^. First, we deconvolute the bulk RNA-seq data into immune cell-type fractions that changed as the patients went from the initial hospitalization through recovery. We show that the estimated values of the immune deconvolution methods approximate the CBC information. We further elucidate that with computational immune deconvolution methods, we can reveal changes between patients infected with SARS-CoV-2 variants with differing severity of disease^48,49^. Next, we illustrate how we can utilize BCR and TCR computational analysis to classify the patients’ cause of disease and investigate the effect of sequencing at various depths on the robustness of our results.

### Approximated immune cell abundances by immune deconvolution methods are close to real complete blood count data

In Figure 1, we observe a consistent positive correlation across lymphocytes, monocytes, and neutrophils using the four deconvolution methods with the CBC data for patients with the Alpha and Alpha+EK (Alpha with an additional E484K mutation at the spike protein) variant infections. The strength of these correlations varies as scores fluctuate in magnitude based on the cell type and method employed. The chart highlights that the EPIC method’s outcomes closely align with the CBC data, standing out, particularly in its accuracy for monocytes. While both quanTIseq and MCP-counter yield commendable results for neutrophils and lymphocytes, xCell’s predictions for neutrophils appear to be less reliable. Importantly, when consolidating the findings from all methodologies, the immune deconvolution results consistently align with the CBC data. However, neutrophils and monocytes especially appear to be harder to estimate overall using deconvolution. Method choice can play an important role here, as only EPIC is able to detect monocytes consistently. This reaffirms that immune deconvolution could serve as an instrument for assessing immune cell levels derived from RNA-seq data, though results do vary between methods and cell types.

**Figure 1:**
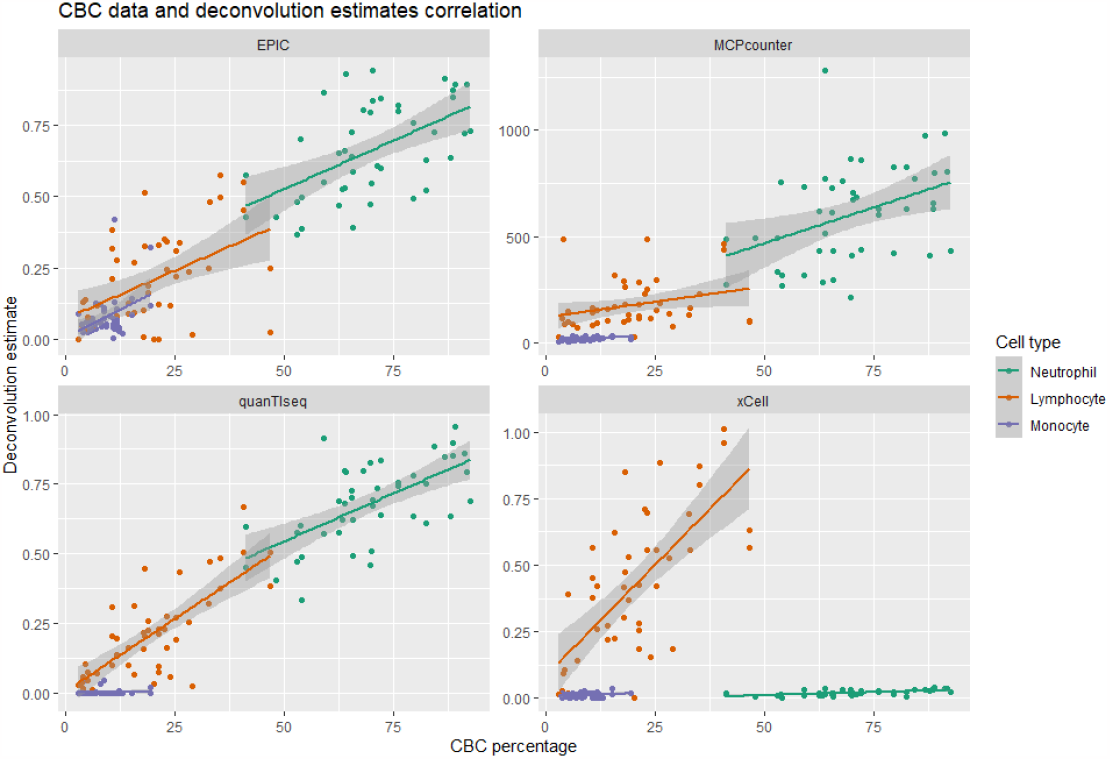
Correlation between CBC data and immune deconvolution scores across lymphocytes, monocytes, and neutrophils using all four deconvolution methods. The colored lines display the linear regression model for each cell type, the shaded areas are the confidence intervals. *Statistical values: EPIC: Neutrophil: R=0*.*55, p=0*.*00011, RMSE=70*.*39, Lymphocyte: R=0*.*47, p=9e-04, RMSE=22*.*61, Monocyte: R=0*.*41, p=0*.*0058, RMSE=9*.*69; MCPcounter: Neutrophil: R=0*.*4, p=0*.*0074, RMSE=576*.*76, Lymphocyte: R=0*.*29, p=0*.*051, RMSE=193*.*25, Monocyte: R=0*.*49, p=0*.*00068, RMSE=13*.*29; quanTIseq: Neutrophil: R=0*.*63, p=4*.*5e-06, RMSE=70*.*37, Lymphocyte: R=0*.*75, p=1*.*8e-09, RMSE=22*.*58, Monocyte: R=-0*.*029, p=0*.*85, RMSE=9*.*77; xCell: Neutrophil: R=0*.*62, p=7*.*5e-06, RMSE=71*.*03, Lymphocyte: R=0*.*69, p=9*.*6e-08, RMSE=22*.*37, Monocyte: R=0*.*24, p=0*.*11, RMSE=9*.*76*.

### Immune deconvolution revealed differences in patients with different severity of disease progression

During the SARS-CoV-2 pandemic, different SARS-CoV-2 variants emerged (ancestral, Alpha, Alpha+EK, Gamma, Omikron BA.1, and Omikron BA.2), which differed in transmissibility and severity. Variants that emerged during the end of the pandemic were associated with less severe disease^50^. The Alpha variant was reported to demonstrate an increase in transmissibility due to the N501Y mutation in comparison to the wild-type virus^51^. The Alpha+EK variant was reported to more efficiently evade a neutralizing antibody response due to the additional E484K mutation but was not associated with more severe disease. The Gamma variant carried both N501Y and E484K mutations and was reported to enhance transmissibility with potential antibody resistance, but, again, disease severity was reported unchanged. The Omicron BA.1 and BA.2 variants, with numerous spike protein mutations, mediated immune escape. However, while their transmissibility increased, these variants, in general, demonstrated less severe disease outcomes. This pattern of viral evolution has been reported previously as a virus adapts to its human hosts over time, favoring transmission over severity^50^.

We hypothesize that non-hospitalized patients infected with an Omicron variant might demonstrate an immune response closer to healthy, non-infected individuals than to hospitalized patients infected with earlier SARS-CoV-2 variants. To explore this hypothesis, we first compared trends in the abundance of B cells, Neutrophils, T cell CD4+, and T cell CD8+ in the SARS-CoV-2 variants and the seronegative samples using four different deconvolution tools (quanTIseq, MCP-counter, EPIC, and xCell, see Materials and Methods) (Figure 2 and Figure 3). We found that all four methods, in general, recapitulated the same trends. The immune cell fractions or scores across all methods and immune cell types evolved over time to more closely resemble seronegative samples as healthy patients. We also observed that non-hospitalized patients infected with an Omicron variant more closely resembled the seronegative patients as compared to the hospitalized patients infected with earlier variants, especially as compared to the time when they were initially hospitalized.

**Figure 2.**
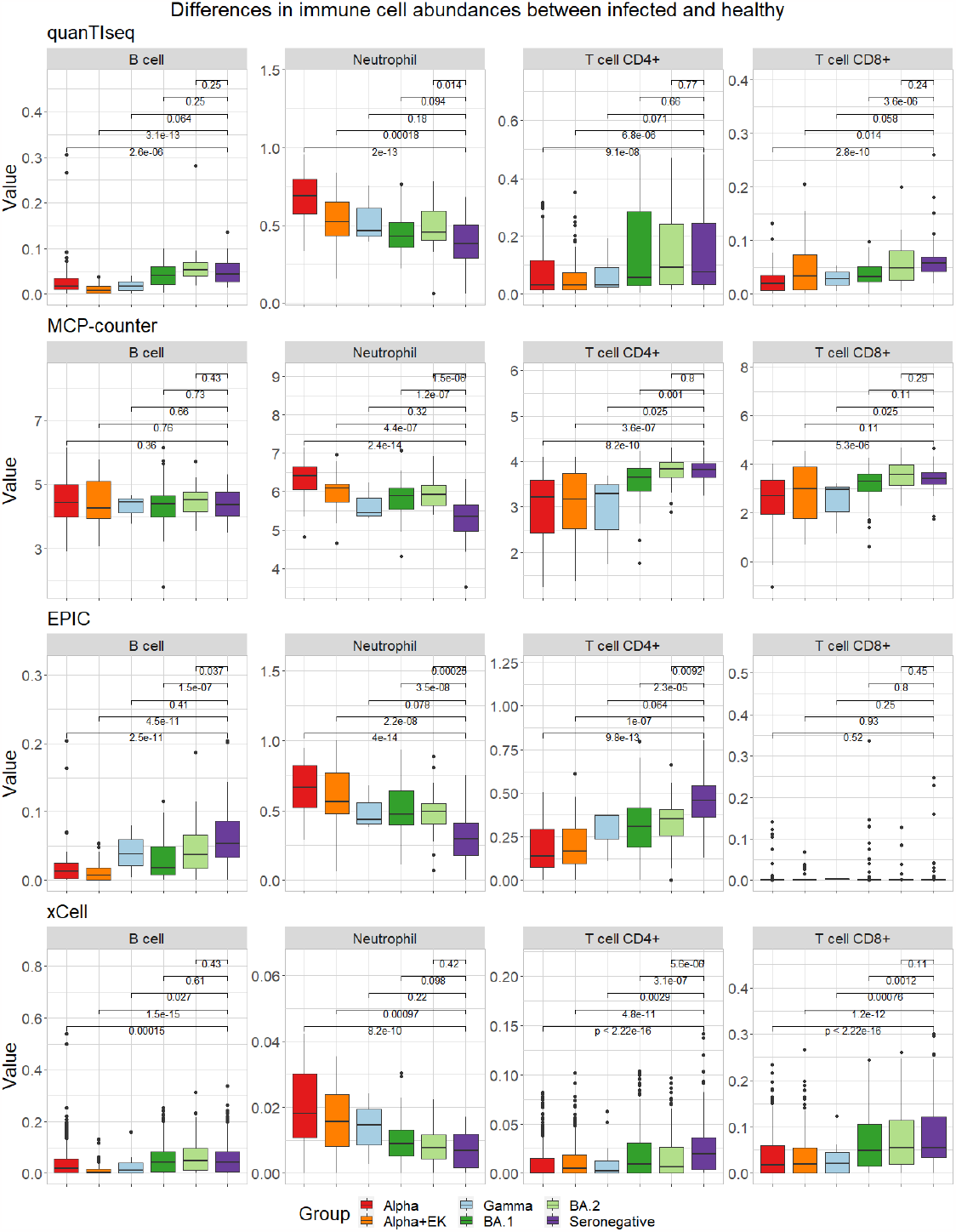
The abundance of immune cells (given by percentage or method-specific score) detected by the immune deconvolution methods quanTIseq, MCP-counter, EPIC, and xCell over all time points combined for the immune cells B cell, Neutrophil, T cell CD4+, and T cell CD8+.

**Figure 3:**
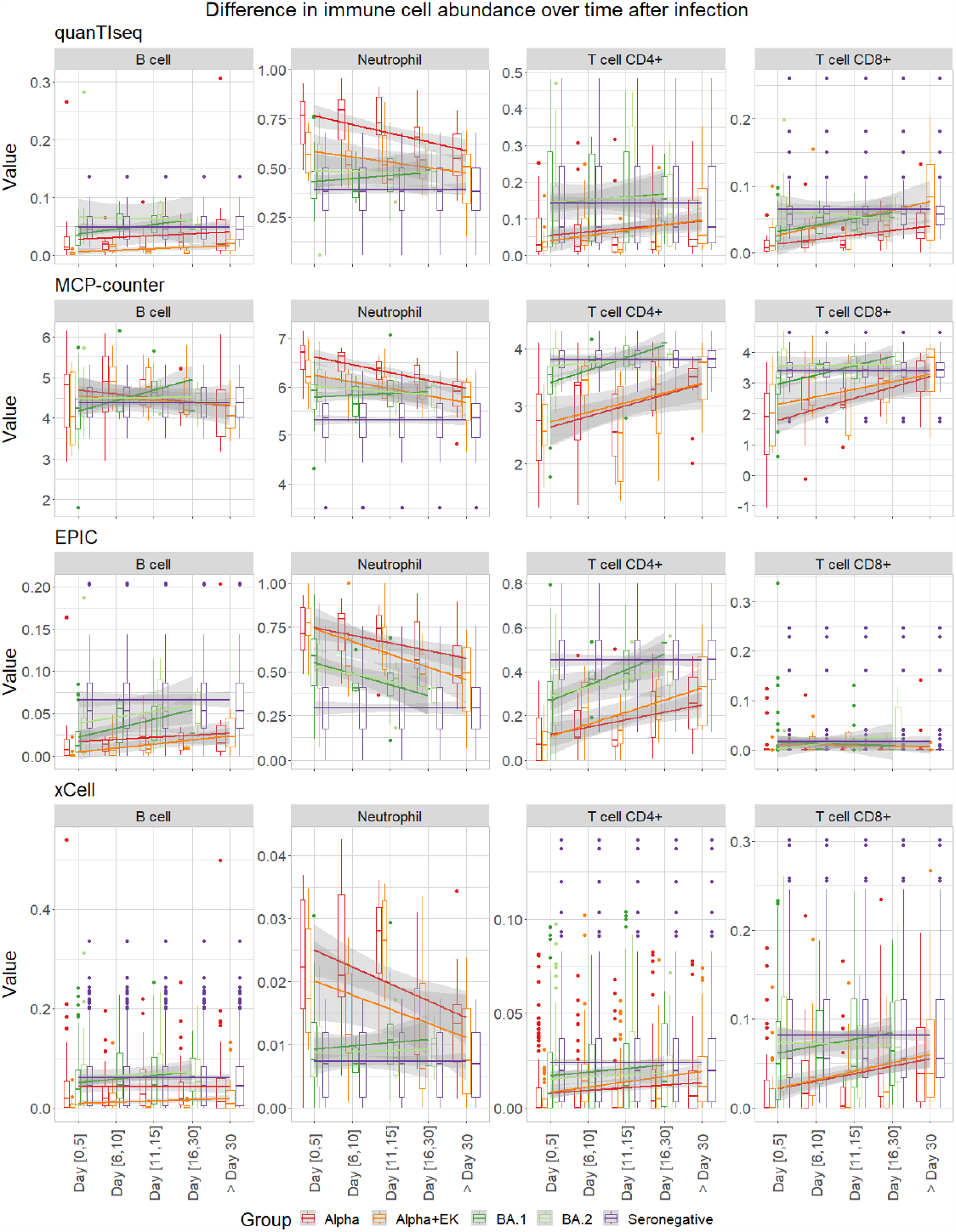
Cell-type fractions separated over brackets 0-5, 6-10, 11-15, 16-30, and >30 days after hospitalization or onset of symptoms detected by the immune deconvolution methods quanTseq, MCP-counter, EPIC, and xCell for the immune cells B cell, Neutrophil, T cell CD4+, and T cell CD8+. The Gamma variant has been removed in this analysis due to poor sample size per time bracket (Suppl. Tables 1a-b).

We further categorized samples into different time brackets after hospitalization or onset of symptoms (days 0-5, 6-10, 11-15, 16-30, and >30). Over time, the projected immune cell fractions appeared to progressively align with those observed in seronegative samples, consistent with patient recovery over time (Figure 3). Patients diagnosed with Alpha and Alpha+EK, variants associated with more severe disease, demonstrated a lengthier time until their immune cell fractions approximated those of seronegative individuals.

### B cell and T cell repertoire analysis offer insights into past or current infections

In general, when an infection occurs, an ‘immunological footprint’ in the form of specific BCR and TCR repertoires can be identified. In this section, we investigated whether bioinformatics BCR and TCR repertoire analysis approaches (i.e., a combination of MiXCR and TRUST4) of transcriptomic data, coupled with a computational tool that associates known BCR and TCR repertoires with causes of diseases (i.e., BLASTp^44,45^), could be used to classify a disease cause for an admitted patient (see Materials and Methods).

With the computational tool MiXCR, we identified 534 unique receptor sequences, while we identified 569 sequencing with TRUST4 across the variants. Of these, 492 sequences were identified by both tools, while 42 and 77 sequences were uniquely identified by MiXCR and TRUST4, respectively. This means that 81% of the sequences were found by both tools, 7% only by MiXCR, and 13% only by TRUST4 (Suppl. Fig. 2). We decided to use only the sequences identified by both tools for further analyses to ensure more reliable results. In the next step, we eliminated sequences that exhibited homology to seronegative samples to account for BCR and TCR sequences that probably lack specificity for SARS-CoV-2, given that no seronegative sample should possess them (see Materials and Methods, Suppl. Fig. 3). Among the residual sequences, we discerned fifteen that did not display similarity to any sequence also found in seronegative samples. A subsequent BLASTp assessment of these sequences identified anti-SARS-CoV-2 immunoglobulin hits within the top 100 matches for seven sequences (Suppl. Table 3). The residual eight sequences predominantly align with generic immunoglobulin sequences. The sequence logo derived from all fifteen sequences highlights conserved motifs at the start (S), the end (VF), and a recurring pattern (DSS) in the center. In contrast, the intervening positions exhibit significant variability, underscoring the pronounced diversity among these sequences (see Suppl. Fig. 4).

### Sequencing depth analysis reveals that a low depth of 10 million reads is sufficient to get to the same conclusions

Despite a reduction in sequencing depth, the trends observed in immune deconvolution outcomes remained consistent. Notably, there were still significant discrepancies in the levels of immune cells when comparing Alpha and Alpha+EK infections to seronegative cases with a sequencing depth of 50 million (Suppl. Fig. 5a) and with an even lower sequencing depth of 10 million (Suppl. Fig. 5b). Furthermore, temporal analysis reaffirmed these findings, indicating the recurrent trend where, across all variants, there is a convergence toward the levels observed in seronegative samples (Suppl. Figs. 6a,b). However, conducting immune deconvolution analysis at a lower sequencing depth results in findings of less significance and lower confidence. Still, the differences described remain significant. Through this analysis, we demonstrated that a lower and even a very low sequencing depth of 10 million is indeed sufficient to discern the trends in immune cell levels and highlight the differential impacts of various variants on the immune system (Figure 4). Furthermore, when evaluating the immune deconvolution scores alongside the CBC data for patients infected with the Alpha and Alpha+EK variants, a notable correlation is evident for a sequencing depth of 50 million (*P*-value 3.765e-11, Suppl. Fig. 7a) and a sequencing depth of 10 million (*P*-value 9.119e-12, Suppl. Fig. 7b).

**Figure 4:**
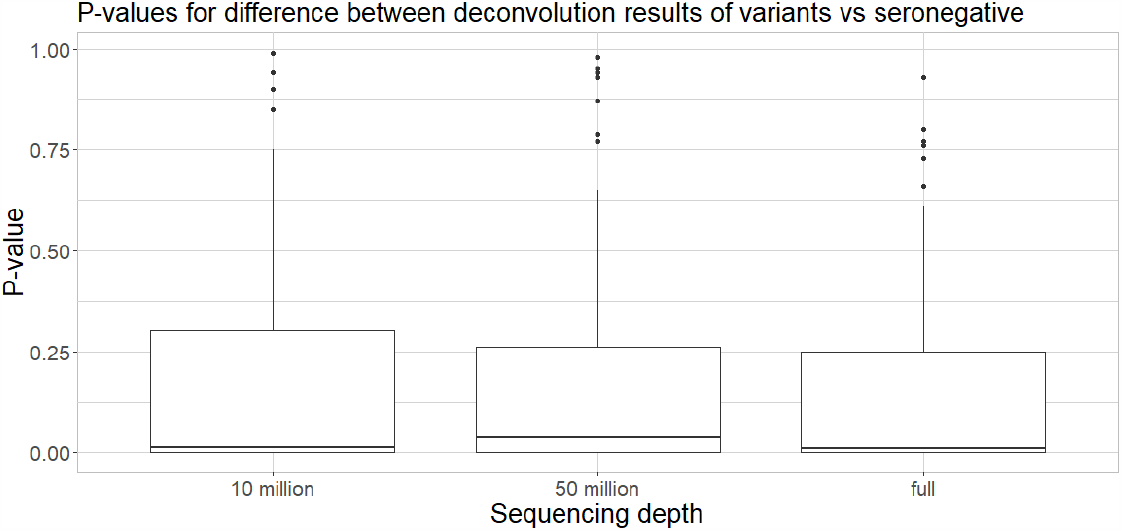
Significance values for sequencing depth analysis for downsampled (10 million and 50 million reads per sample) and the full sequencing depth (up to 150 million reads per sample) in the immune deconvolution analysis.

In the repeated BCR/TCR analysis with a significantly reduced sequencing depth of 10 million, we identified only 95 unique BCR and TCR sequences. This is markedly fewer than in the prior analysis, but the decline is anticipated due to the reduced sequencing depth, which results in fewer overall sequences from the RNAseq experiments. Of the 95 sequences, 18 (19%) were solely identified by MiXCR, seven (7%) exclusively by TRUST4, and 70 (74%) were detected by both tools. This indicates that the majority of the sequences were still identifiable by both tools (Suppl. Fig. 8). After eliminating sequences resembling those in seronegative samples, we pinpointed eight unique sequences. Among these, seven were matched to anti-SARS-CoV-2 immunoglobulin sequences (Suppl. Table 4, Suppl. Figs. 7 and 8).

The sequences identified by the two BCR/TCR analyses, with full sequencing depth, and with low sequencing depth, differ between results. Additionally, there’s a variation in the positions of the SARS-CoV-2 specific hits. At greater sequencing depth, these hits are more commonly found within the top ten. In contrast, when the sequencing depth is reduced, they are more likely to be ranked higher, and, as a result, the findings become somewhat less substantiated.

A statistical comparison like comparing the *P*-values for the immune deconvolution is not possible here as MiXCR and TRUST4 do not generate significance values, and the BLAST *E*-values represent the number of random hits that can be generated in a database of a certain size and, therefore are not suitable to compare the significance of our results but merely the reliability of each sequence match individually. In both analyses, we were able to find seven anti-SARS-CoV-2-related hits that appear in the first one hundred BLAST results. Notably, even though MiXCR and TRUST4 identified fewer sequences overall due to the reduced depth, the count of SARS-CoV-2 specific sequences remained consistent.

In conclusion, a sequencing depth of 10 million was adequate to yield results comparable to analyses with much higher depths (up to 200 million reads). However, a greater sequencing depth produces more robust outcomes.

## Discussion

We found that the immune deconvolution tools, including quanTIseq, MCP-counter, EPIC, and xCell, generally predict similar trends in immune cell composition (B cells, Neutrophils, T cell CD4+, and T cell CD8+) across SARS-CoV-2 samples that reflect differences in severity and over time. However, we can also see large differences between individual samples. Our computational results predict a progressive alignment of immune cell fractions with those of seronegative samples, correlating with decreased disease severity and/or individual disease progression. However, individuals with severe disease courses like Alpha and Alpha+EK show extended recovery timelines before reaching these levels, indicating a potential marker of disease severityA confounder that should be considered in the analysis could be that SARS-CoV-2 can invade immune cells and could potentially skew the results of the immune deconvolution results^52^. While computational deconvolution methods are able to robustly estimate trends in immune-cell composition correctly, they do show a large variance in prediction accuracy on a sample level. This drawback is especially important when trying to use such methods in a personalized fashion. Here, prediction accuracy is not high enough to give precise results of immune-cell composition in patients. However, so-called second-generation deconvolution methods^53^ promise to increase prediction quality by employing scRNA-seq datasets as an additional resource in deciphering the cell-type composition of bulk RNA-seq datasets. Such tools may also reveal changes in the functional state of immune cells and thus surpass information provided by CBC measurements.

We further introduced an approach for diagnosing infections using RNA-seq with bioinformatic analysis of BCR and TCR repertoires. We speculate that patterns of BCR and TCR repertoires could be associated with different disease settings. The current system is built on known BCR and TCR repertoires associated with diseases, which means it can only be used for identifying known infections^54,55^. As data on BCR and TCR repertoires from different clinical settings is deposited and available for analysis, it is possible the information can be used to improve understanding of immune response in individual patients. At present, ethical considerations of detailed genomic analysis in individual patients can limit the types of information gathered and their distribution. However, anonymized data obtained through clinical trials with informed consent may still be useful in exploring how changes in TCR and BCR repertoires evolve during disease and recovery.

Our analyses demonstrate that a reduced sequencing depth of 10 million is sufficient to identify overarching trends in immune cell levels and anti-SARS-CoV-2 specific sequences, although higher sequencing depths yield more robust outcomes. Despite lower depths resulting in findings of less significance and confidence, the overall trends and correlations with CBC data remain consistent. The BCR/TCR analyses further corroborate these findings, as even at reduced sequencing depths, SARS-CoV-2-specific sequences were still identifiable. These results affirm the feasibility of using lower sequencing depths for meaningful analyses in the study of immune responses and pathogen-specific immunity, making it more feasible in a clinical setting due to lower costs.

Since 2001, genome sequencing costs have significantly decreased from $100 million to the $1,000 genome milestone, reflecting similar cost reductions in RNA sequencing^56^. With the impending expiration of Illumina’s key patents, the RNA sequencing market could see heightened competition and further price reductions, a recent article in Science just speculated about the costs being reduced to $100^57^. This shift might be key to embedding sequencing more deeply into routine clinical practice, making it a more accessible tool for patient care and research.

As RNA-sequencing technologies advance and become cheaper, they hold promise for future clinical utility by providing a more detailed view of global gene expression profiles. For example, quantitative polymerase chain reaction (qPCR) has already been adopted in clinical settings for its high sensitivity and specificity in detecting and quantifying microbial pathogens^58^ or SARS-CoV-2^59,60^. To our knowledge, RNA-seq combined with immune deconvolution is not directly used in a routine clinical setting^61^; however, it has been employed in research settings analyzing whole blood sequencing datasets^14^. Previous work has also identified specific immune cell subsets, including neutrophils, to be associated with more severe SARS-CoV2 infection^62^. In the future, immune deconvolution and BCR/TCR could potentially guide the decision-making of a physician, e.g., the immediate allocation of a newly admitted patient with potentially severe disease progression to the intensive care unit, recognizing that such a tool would require ongoing updates to maintain its utility for predictive modeling^63,64^. With more blood samplings after admission, we can also see if the disease course will change, and the medical doctor could, based on this analysis and other factors, advise the patient to be submitted to the intensive care unit^65^. Nonetheless, we have to consider that our study relies on data either primarily collected from hospitalized elderly patients (i.e., Alpha, Alpha+EK, and Gamma) or mild disease progression (i.e., Omikron BA.1 and Omikron BA.2), potentially introducing a selection bias. Moreover, differences in local healthcare systems, as well as individual patient factors (e.g., age and preconditions), could influence recovery timelines and should be factored into any broader applications of these findings. Additionally, the predictive methods used for immune cell fraction estimations, while robust and consistent, are not without their limitations and potential discrepancies. In addition to immune cell composition, analyzing immune cell receptors of B and T cells by employing tools such as MiXCR^33^ and TRUST4^34^ in combination with BCR/TCR databases can provide a rapid determination of the type of a previously discovered virus or infection. However, with our proposed method, we do find potential clonotypes but are not able to confirm if they come from a new virus variant. Genomic data analysis from cell preparation, library generation, sequencing, and quality control is, with the current technology, not feasible in a matter of hours, as is the case for CBCs. Recent advances to introduce RNA-seq into clinical settings describe a complete workflow to finish in about one week^66^. One technology that is able to improve the precision of BCR/TCR detection is Oxford Nanopore sequencing. While not currently implemented in many studies of the transcriptome due to sequencing error limitations and PCR-induced distortions^67^, it promises to increase clonotype detection and tracking^68^.

In summary, we employed computational immune deconvolution tools at distinct SARS-CoV-2 data sets, illustrating that they can be used to supplement immune cell abundance estimates for bulk RNA-seq data that is not accompanied by CBC information. Additionally, these tools can be used for discerning trends in immune cell fractions during disease recovery and for comparing differences in immune cell fractions between more and less severe SARS-CoV-2 variants. Using the proposed workflow to utilize BCR/TCR methods combined with alignments and pBLAST could help to pinpoint the type of viral infection. Our presented bioinformatic strategies combined with expert medical judgment, new technologies, and automatizations could promise a path toward precision medicine, where treatment plans are personalized and optimized for each individual in the future based on individualized genetic analyses.

## Supporting information

Supplementary Data is available at biorxiv online.

## Availability

*Computational scripts can be found at:* https://github.com/biomedbigdata/SARS-CoV-2_immunedeconv_bcrtcr

*Analysis results can be downloaded as an RData object in Supplemental Materials and on figshare:* www.doi.org/10.6084/m9.figshare.24221167

*Data can be publicly found at:*

GSE190680 (variants: Alpha, Alpha+EK, Gamma) GSE162562 (Seronegatives)

GSE201530 (variant: Omikron BA.1)

GSE205244 (variants: Omikron BA.1 and Omikron BA.2)

List of marker genes per method can be found here: https://doi.org/10.6084/m9.figshare.24442423.v1

## Acknowledgments

The authors gratefully thank all the patients and healthy individuals who participated in this study. We would like to thank Anke Kraft and Sebastian Klein for their helpful discussions. Figures were created with Biorender.com. Parts of the figures include icons from Flaticon.com under a paid license. The text was partly rephrased using chatGPT version 4 under a paid license.

## Authorship Contributions

MH, LLW, AD, HKL, LK, PAF, LH, and ML contributed to the initial design of the study. MH and LLW conducted the data cleaning and analyses. MD and AD provided supervision of LLW during the analyses. NT supported data cleaning, data visualization, and technical work. MH, LLW, and AD drafted the initial manuscript. MH, LLW, AD, MH, PAF, LH, and ML edited the initial manuscript. All authors read and approved the final version of the manuscript.

