## Supplementary Data is available at biorxiv online. for "Blood transcriptomics analysis offers insights into variant-specific immune response to SARS-CoV-2"

[1] Big Data in BioMedicine Group, Chair of Experimental Bioinformatics, TUM School of Life Sciences, Technical University of Munich, Munich, Germany
[2] Institute for Advanced Study (Lichtenbergstrasse 2 a, D-85748 Garching, Germany), Technical University of Munich, Germany
[3] National Institute of Diabetes, Digestive, and Kidney Diseases, Bethesda, MD 20892, United States of America
[4] TyrolPath Obrist Brunhuber GMBH, Zams, Austria
[5] Chair of Computational Systems Biology, University of Hamburg, Hamburg, Germany
[6] Computational BioMedicine Lab, University of Southern Denmark, Odense, Denmark
[7] Departments of Oncology & Medicine, Georgetown University, Washington, DC, United States of America
*The authors wish it to be known that, in their opinion, the first three authors should be regarded as Joint First Authors.

### ABSTRACT

Bulk RNA sequencing (RNA-seq) of blood is typically used for gene expression analysis in biomedical research but is still rarely used in clinical practice. In this study, we argue that RNA-seq should be considered a routine diagnostic tool, as it offers not only insights into aberrant gene expression and splicing but also delivers additional readouts on immune cell type composition as well as B-cell and T-cell receptor (BCR/TCR) repertoires. We demonstrate that RNA-seq offers vital insights into a patient's immune status via integrative analysis of RNA-seq data from patients infected with various SARS-CoV-2 variants (in total 240 samples with up to 200 million reads sequencing depth). We compare the results of computational cell-type deconvolution methods (e.g., MCP-counter, xCell, EPIC, quanTIseq) to complete blood count data, the current gold standard in clinical practice. We observed varying levels of lymphocyte depletion and significant differences in neutrophil levels between SARS-CoV-2 variants. Additionally, we identify B and T cell receptor (BCR/TCR) sequences using the tools MiXCR and TRUST4 to show that - combined with sequence alignments and pBLAST - they could be used to classify a patient's disease. Finally, we investigated the sequencing depth required for such analyses and concluded that 10 million reads per sample is sufficient. In conclusion, our study reveals that computational cell-type deconvolution and BCR/TCR methods using bulk RNA-seq analyses can supplement missing CBC data and offer insights into immune responses, disease severity, and pathogen-specific immunity, all achievable with a sequencing depth of 10 million reads per sample.

### Supplemental Materials 1: Immune deconvolution methodology

MCP-counter and xCell are marker-gene-based cell-type deconvolution methods that leverage cell-type-specific marker genes to compute an enrichment score that can be compared across samples. Using gene expression data as input, MCP-counter computes scores for ten different cell types[1], whereas xCell computes scores for 64 different cell types[2].

The other two methods we use, EPIC[3] and quanTIseq[4], consider the deconvolution problem as a set of equations based on $b = C \times p$ (1), where $b$ represents the gene expression matrix, $C$ is a signature matrix containing a reference gene expression profile for all cell types and $p$, the vector denoting the proportions of each cell type, which are estimated. Both methods come with precomputed signature matrices within the immunedeconv package.

Each of the four methods gives results for various cell types. For example, xCell identifies 64 specialized cell types, while EPIC focuses on just six broader categories. To compare them fairly, we used a provided cell-type mapping in the immunedeconv package that translates the cell types into a standard language, which is called "controlled vocabulary cell types" that map specific cell types to broader cell types, e.g., mapping regulatory T cells to CD4+ T cells and allow a side-by-side comparison across methods.

### Supplemental Materials 2: BCR/TCR repertoire methodology

MiXCR aligns sample reads to a customized database from GeneBank[5], groups them into clonotypes (i.e., receptor sequences that target the same antigen), corrects for polymerase chain reaction and sequencing errors, and exports the results in a tab-delimited file. TRUST4, in contrast, isolates candidate reads and carries out de novo assembly (i.e., this means it constructs longer sequences, or "contigs", from short reads based solely on their overlapping segments, without relying on any reference sequence). Then, it annotates the assembled contigs by aligning them to the ImmunoGeneTics database[6] and reports matched CDR3 sequences (i.e., unique areas of the antigen receptor gene that play a crucial role in the diversity of the immune response by determining the specific antigen-binding affinity of T and B cell receptors[7]).

### Supplemental Materials 3: ClustalW algorithm in the BCR/TCR repertoire methodology

We use the ClustalW algorithm[8] to perform multiple sequence alignment via the *calc_dist_mat()* method from the AlignmentDistanceCalculator class in scirpy and the BLOSUM62[9] similarity matrix. Using the R package igraph[10] and the computed distances, we constructed a graph with sequences as nodes and their similarities represented by edges. This graph allowed visualization of method overlap by coloring nodes based on the reconstructing method that identified the sequence (either one of MiXCR and TRUST4 or both) and pinpointed unique sequences specific to SARS-CoV-2 variants by spotlighting sequences exclusive to infected samples and without connection to sequences from seronegative samples. According to the scirpy documentation, a sensible sequence distance cutoff is below 15, though we adopted a stricter threshold of 10, suggesting CDR3 sequences below this limit likely target the same antigen. For further analysis, we generated multiple sequence alignments of the reconstructed BCR and TCR CDR3 sequences using the msa R package[11] and depicted sequence conservation and regions with higher variation through sequence logos crafted with the ggseqlogo R package[12].

**Supplemental Table 1: Overview of the number of samples**

| Variant \ time | N/A | Day 0-5 | Day 6-10 | Day 11-15 | Day 16-30 | > Day 30 | total |
|---|---|---|---|---|---|---|---|
| Alpha | - | 18 | 12 | 10 | 17 | 10 | 67 |
| Alpha EK | - | 8 | 7 | 4 | 4 | 7 | 30 |
| Gamma | - | 1 | 1 | 1 | - | - | 3 |
| Omicron BA.1 | - | 50 | 5 | 23 | 1 | - | 79 |
| Omicron BA.2 | - | 13 | - | 5 | 5 | - | 23 |
| Seronegative | 50 | - | - | - | - | - | 50 |
| total | 50 | 90 | 25 | 43 | 27 | 17 | 252 |

**Supplemental Table 1a:** Table of the total number of samples per time point in the GEO database.

| Variant \ time | N/A | Day 0-5 | Day 6-10 | Day 11-15 | Day 16-30 | > Day 30 | total |
|---|---|---|---|---|---|---|---|
| Alpha | - | 16 | 11 | 8 | 17 | 9 | 61 |
| Alpha EK | - | 8 | 6 | 4 | 4 | 7 | 29 |
| Gamma | - | 1 | 1 | 1 | - | - | 3 |
| Omicron BA.1 | - | 49 | 5 | 23 | 1 | - | 78 |
| Omicron BA.2 | - | 12 | - | 5 | 5 | - | 22 |
| Seronegative | 50 | - | - | - | - | - | 47 |
| total | 50 | 86 | 23 | 41 | 47 | 16 | 240 |

**Supplemental Table 1b:** Table of the number of samples per time point used for downstream analysis after quality control.

### Supplemental Table 2: Excluded sample due to insufficient quality

| sample | reason(s) |
|---|---|
| ID43_1st | distance to other samples, distance on PCA plot |
| B.1.351-ID5_3rd | problem with STAR alignment, failed threshold of minimum mapped reads |
| B.1.351-ID1_3rd_2 | problem with GC content, not normally distributed[13] |
| ID29_2nd_2 | problem with GC content, not normally distributed[13] |
| ID34_2nd_2 | problem with GC content, not normally distributed[13] |
| ID38_3rd_2 | problem with GC content, not normally distributed[13] |
| ID53_2nd_2 | problem with GC content, not normally distributed[13] |
| BNT_Aus_20_2 | problem with GC content, not normally distributed, distance to other samples, gene counts distributed badly (peak on chr 21) |
| A_118_Asymptom | distance to other samples, distance on PCA plot |
| B.1.351.ID14_1st | first sampling at day -1 |
| ID_38_2nd | high mitochondrial gene counts (>0.1)[14] |
| ID_38_3rd | high mitochondrial gene counts (>0.1)[14] |
| ID_38_1st | because ID38 has pancytopenia due to myelodysplastic syndrome: decrease in all three peripheral blood cell lines [15] |
| B_425_Seronegative | high mitochondrial gene counts (>0.1)[14] |
| B_436_Seronegative | high mitochondrial gene counts (>0.1)[14] |
| B_446_Seronegative | high mitochondrial gene counts (>0.1)[14] |
| SRR18922948_COVID-19-Omicron_BA-1 | problem with GC content, not normally distributed[13] |
| SRR18922909_COVID-19-Omicron_BA-1 | too many overrepresented sequences, distance to other samples |
| SRR18922902_COVID-19-Omicron_-- | no classification if BA.1 or BA.2 |

**Supplemental Table 2:** List of excluded samples due to quality concerns.

### Supplemental Table 3: Identified BCR and TCR repertoire sequences with no similarities to seronegative BCR and TCR repertoire sequences and their hits with BLASTp in samples up to 200M sequencing depth

| Sequence | Hit | Hit position | E-value |
|---|---|---|---|
| CYSTDSSGNHRGVF | anti-SARS-CoV-2 immunoglobulin lambda [Homo sapiens] | 8 | 6e-05 |
| CNSRDSSGNHLGVF | immunoglobulin light chain junction region [Homo sapiens] | 1 | 1e-05 |
| CQSYDSSNVVF | anti-SARS-CoV-2 immunoglobulin light chain [Homo sapiens] | 7 | 0.050 |
| CMQGTHWPTF | immunoglobulin light chain junction region [Homo sapiens] | 1 | 0.002 |
| CQSYDSSLSGSGVF | immunoglobulin light chain junction region [Homo sapiens] | 1 | 3e-05 |
| CLQHDNFPYTF | immunoglobulin light chain junction region [Homo sapiens] | 1 | 3e-04 |
| CAAWDDSLNGHVVF | immunoglobulin light chain junction region [Homo sapiens] | 1 | 1e-06 |
| CLQHDNFPLTF | (11-NOV-2022) anti-SARS-CoV-2 immunoglobulin light chain [Homo sapiens] [Mus musculus] | 5 | 0.010 |
| CQAWDSSVVF | (15-MAR-2023) anti-SARS-CoV-2 immunoglobulin light chain [Homo sapiens] | 4 | 0.27 |
| CVVSDRGSTLGRLYF | T cell receptor alpha chain V region (clone 1V alpha 24-1) - human (fragment) [Homo sapiens] | 1 | 8e-07 |
| CSSYTSSSTVF | (15-FEB-2023) anti-SARS-CoV-2 immunoglobulin light chain variable region [Homo sapiens] | 82 | 2.6 |
| CQSYDSSLSGSYVF | immunoglobulin light chain junction region [Homo sapiens] | 1 | 7e-06 |
| CQAWDSSTVF | immunoglobulin light chain junction region [Homo sapiens] | 1 | 0.037 |
| CETWDSNTRVF | anti-SARS-CoV-2 spike protein immunoglobulin light chain variable region [Homo sapiens] | 10 | 0.006 |
| CQQRSNWPPTWTF | anti-SARS-CoV-2 immunoglobulin light chain variable region [Homo sapiens] | 8 | 5e-06 |

**Supplemental Table 3:** The fifteen unique sequences that were distinct from healthy BcR and TCR repertoire samples with their first BLASTp result or the first result with SARS-CoV-2 connection, as well as the E-value and position of the result.

### Supplemental Table 4: Identified BCR and TCR repertoire sequences with no similarities to seronegative BCR and TCR repertoire sequences and their hits with BLASTp in samples up to 10M sequencing depth

| Sequence | Hit | Hit position | E-value |
|---|---|---|---|
| CAAWDDSLNGWVF | anti-SARS-CoV-2 immunoglobulin light chain [Homo sapiens] | 61 | 4e-05 |
| CAAWDDSLNGPVF | anti-SARS-CoV-2 immunoglobulin heavy chain variable region [Homo sapiens] | 39 | 2e-04 |
| CQSADSSGTYVVF | anti-SARS-CoV-2 immunoglobulin light chain [Homo sapiens] | 13 | 0.002 |
| CQSYDSSLSGSVF | anti-SARS-CoV-2 immunoglobulin light chain [Homo sapiens] | 34 | 0.003 |
| CGTWDSSLSAGVF | anti-SARS-CoV-2 immunoglobulin light chain [Homo sapiens] | 35 | 0.001 |
| CLQHNSYPWTF | anti-SARS-CoV-2 immunoglobulin light chain variable region [Homo sapiens] | 28 | 0.002 |
| CMQATQFPRTF | anti-SARS-CoV-2 immunoglobulin light chain [Homo sapiens] | 3 | 0.007 |
| CALWEVQELGKKIKVF | T-cell receptor gamma chain [Homo sapiens] | 1 | 7e-09 |

**Supplemental Table 4:** The eight unique sequences that were distinct from healthy BcR and TCR repertoire samples derived from the MiXCR and TRUST4 results for downsampled results with a sequencing depth of 10 Mio. For each sequence, the first BLASTp result or the first result with SARS-CoV-2 connection is shown, as well as the E-value and position of the result.

### Supplemental Figure 1a:

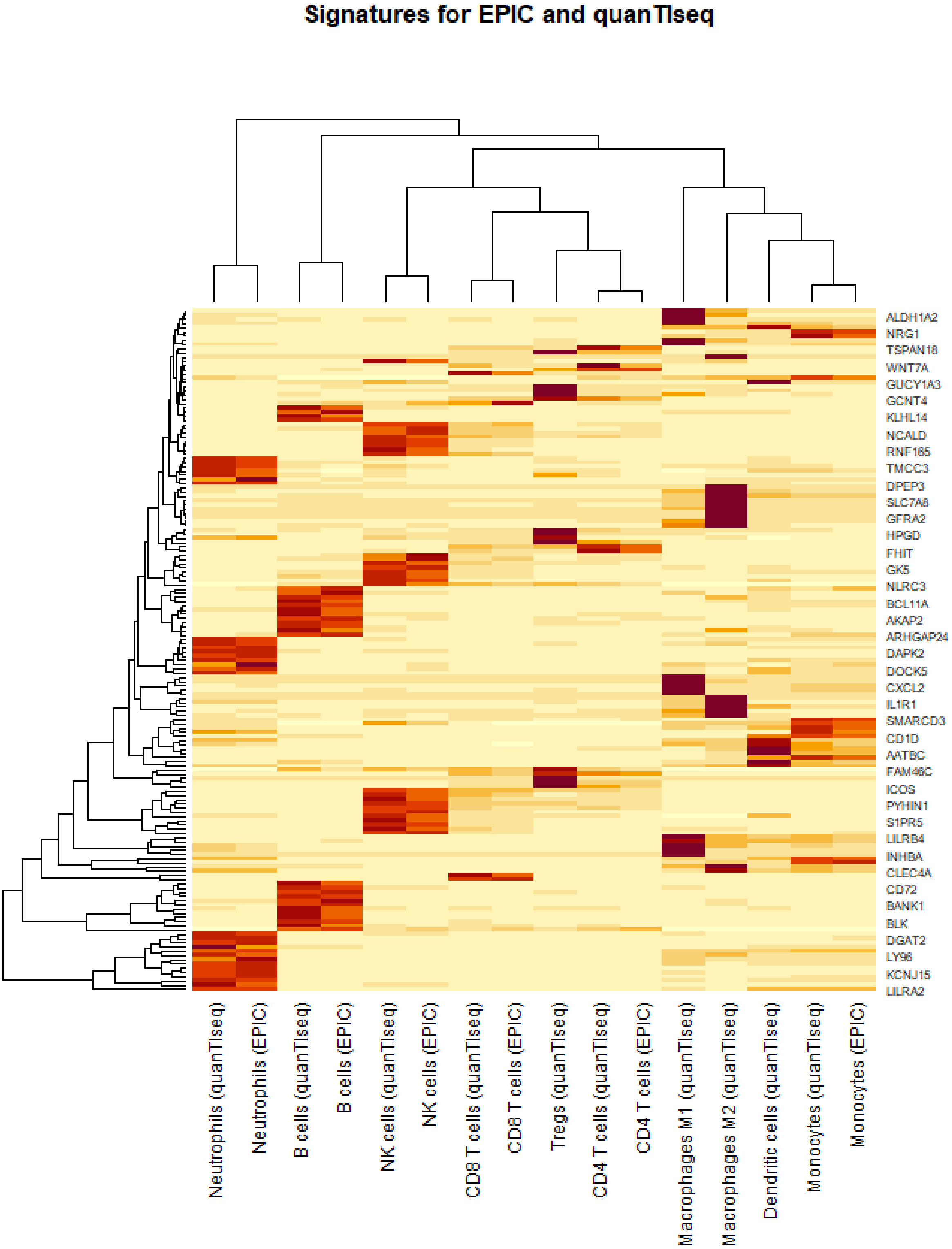


**Supplemental Figure 1a:** We visualized the marker genes shared by quanTIseq and EPIC.

### Supplemental Figure 1b:

**Common transcriptomic markers for MCP-counter and xCell**

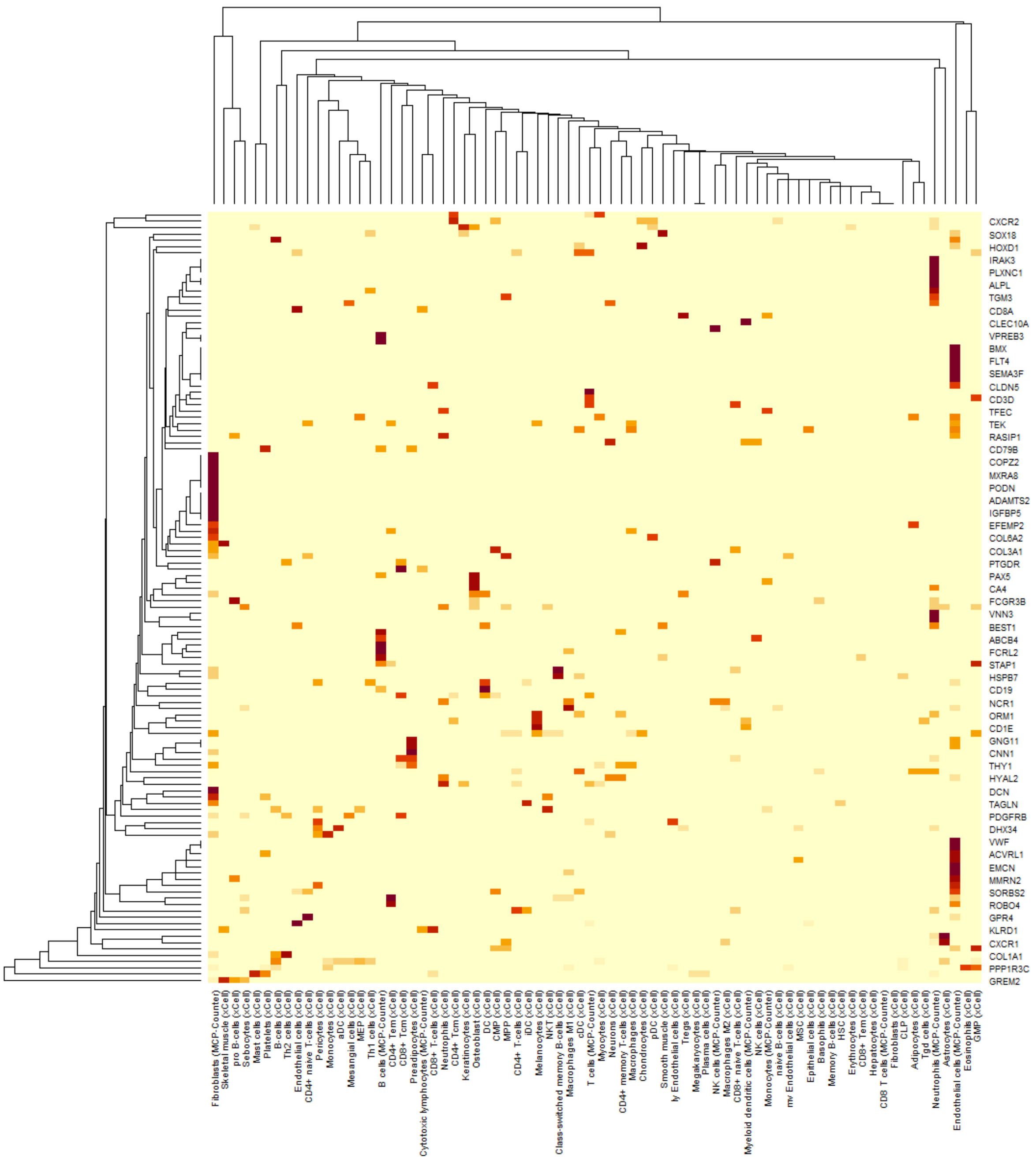


**Supplemental Figure 1a:** We visualized the marker genes shared by MCP-counter and xCell.

### Supplemental Figure 2:

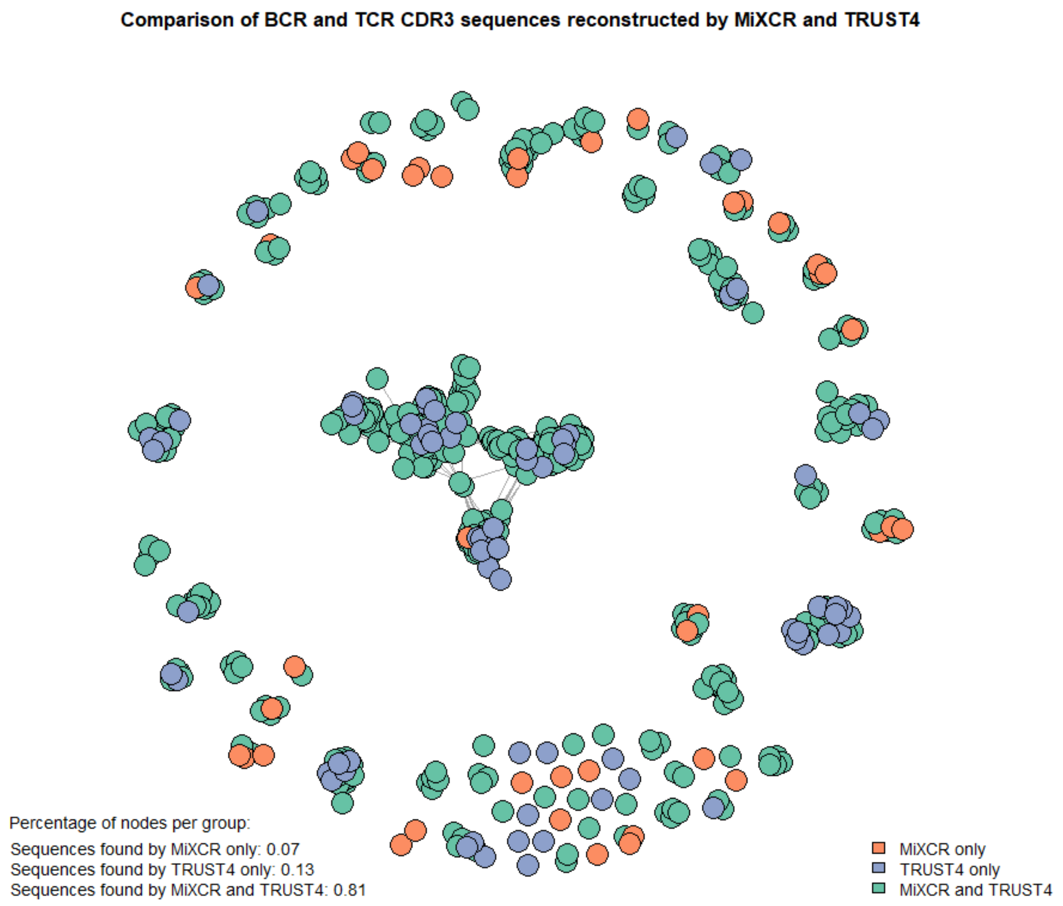


**Supplemental Figure 2:** We visualized the results of MiXCR and TRUST4 in a graph, where sequences are nodes, with an edge between two nodes if they are similar (see Materials and Methods). The sequences found by both tools are represented in green, those found only by MiXCR are in orange, and those found only by TRUST4 are in blue. The majority of nodes are green, confirming that most sequences were found by both tools.

### Supplemental Figure 3:

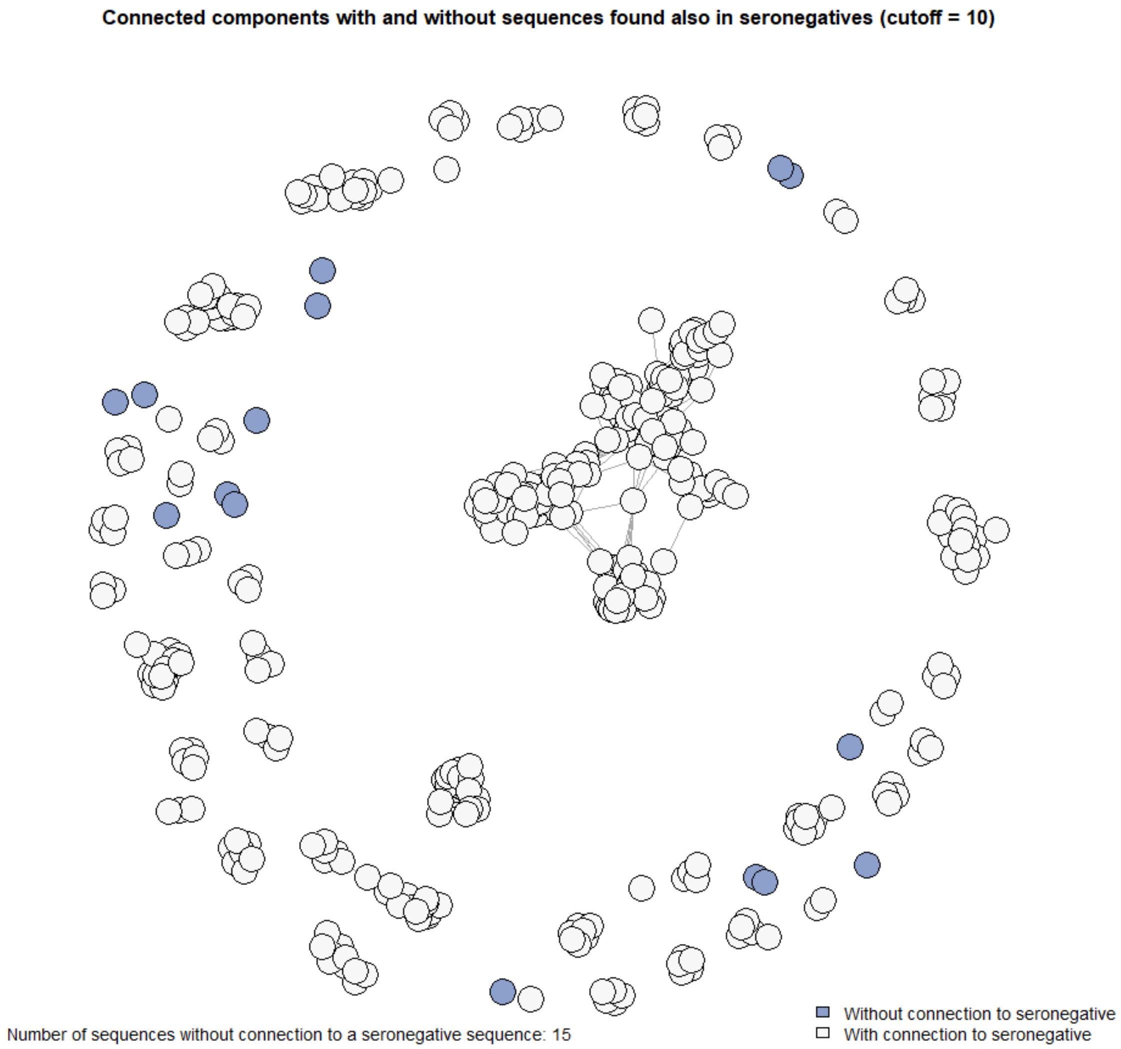


**Supplemental Figure 3:** MiXCR and TRUST4 both predicted 15 BCR/TCR repertoire sequences that were distinct from seronegative samples.

### Supplemental Figure 4:

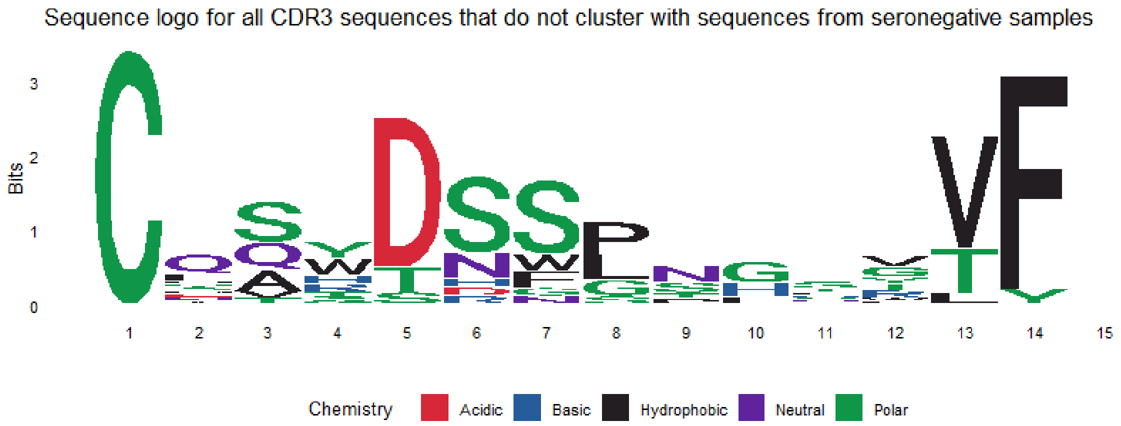


**Supplemental Figure 4:** Sequence logo of the BCR and TCR repertoire sequences that were not similar to the ones found in the Seronegative samples.

**Supplemental Figure 5a:**

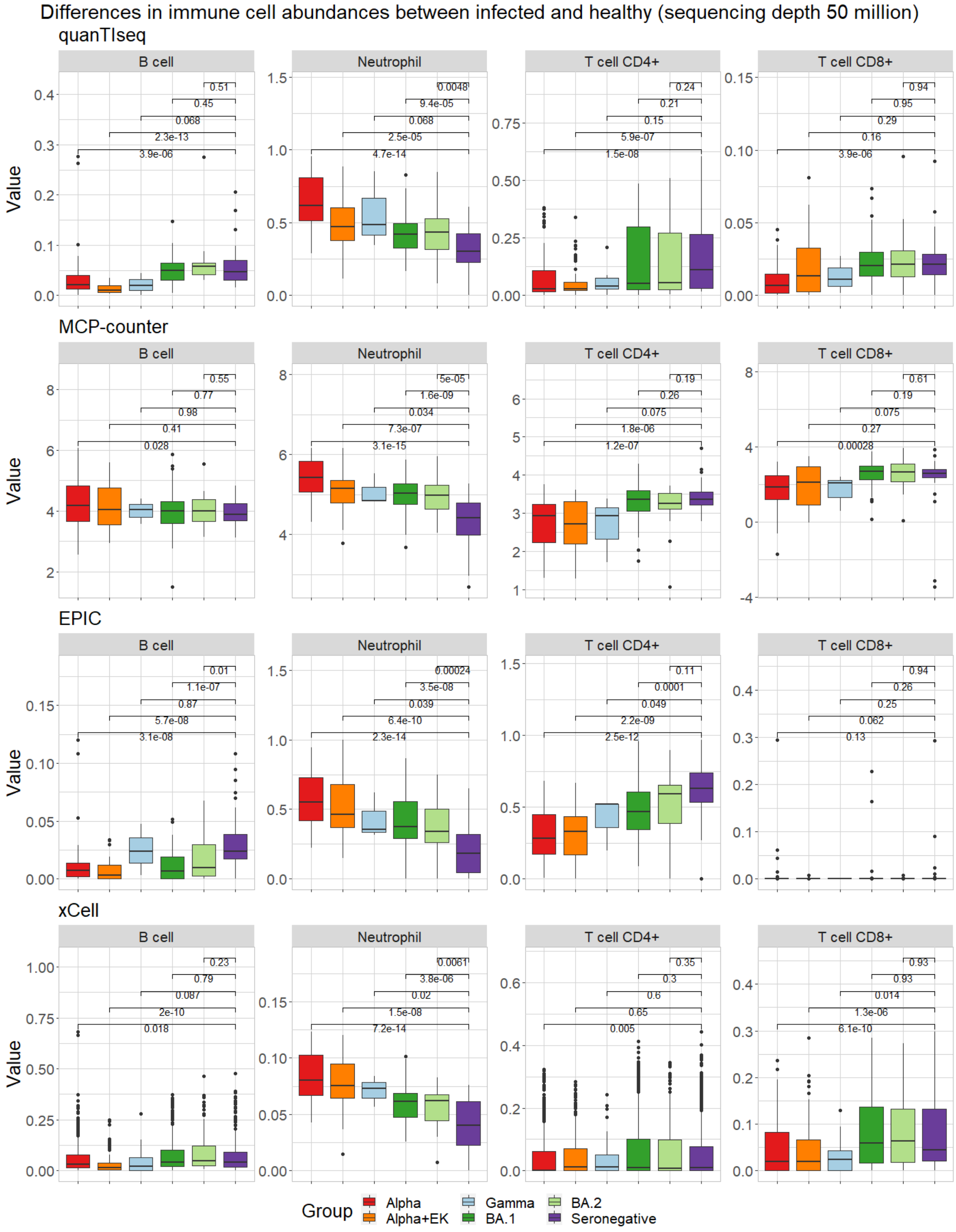


**Supplemental Figure 5a:** Immune deconvolution results for downsampled sequencing depth 50 million. The abundance of immune cells (given by percentage or method-specific score) detected by the immune deconvolution methods quanTIseq, MCP-counter, EPIC, and xCell over all time points combined for the immune cells B cell, Neutrophil, T cell CD4+, and T cell CD8+.

### Supplemental Figure 5b:

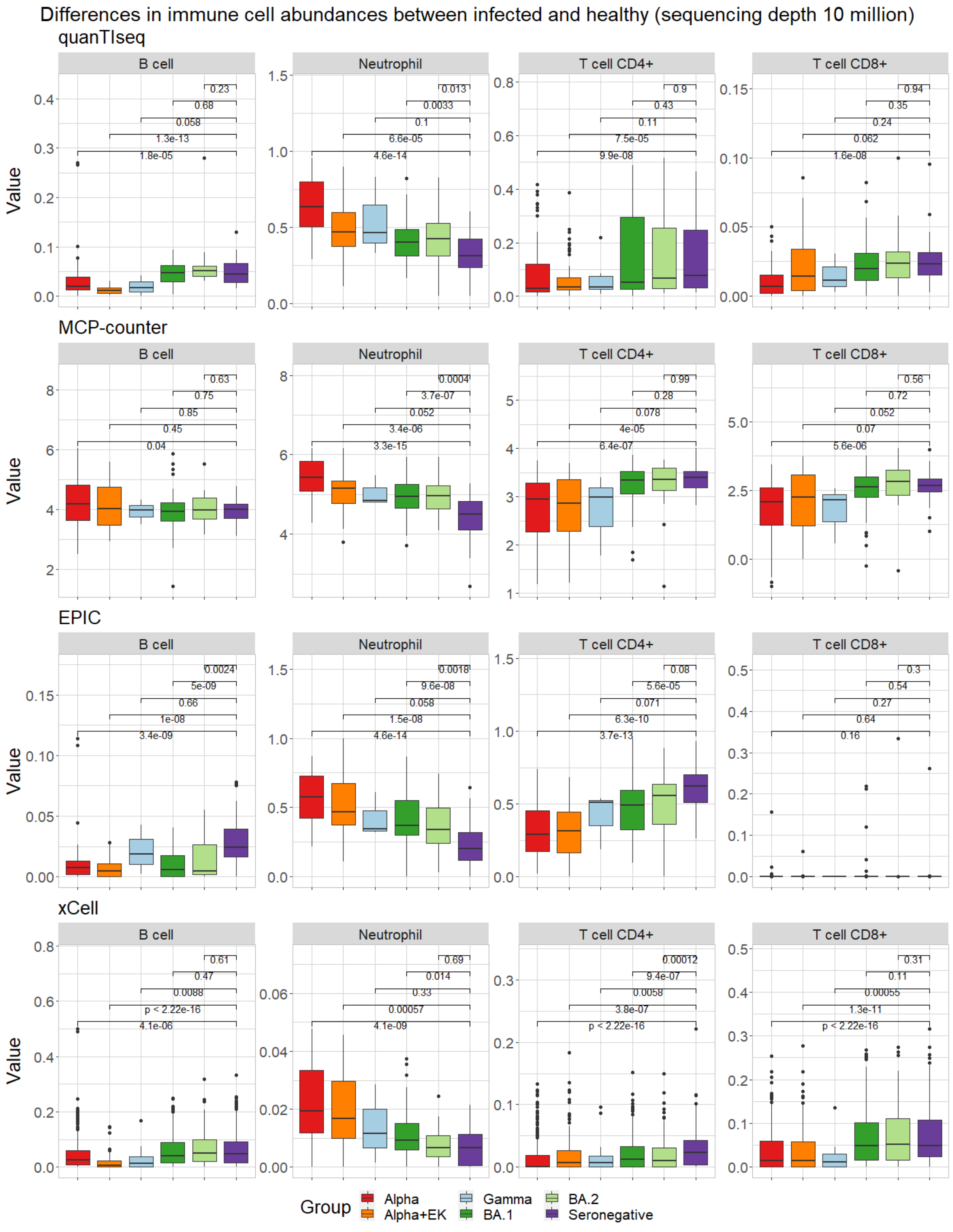


**Supplemental Figure 5b:** Immune deconvolution results for downsampled sequencing depth 10 million. The abundance of immune cells (given by percentage or method-specific score) detected by the immune deconvolution methods quanTIseq, MCP-counter, EPIC, and xCell over all time points combined for the immune cells B cell, Neutrophil, T cell CD4+, and T cell CD8+.

### Supplemental Figure 6a:

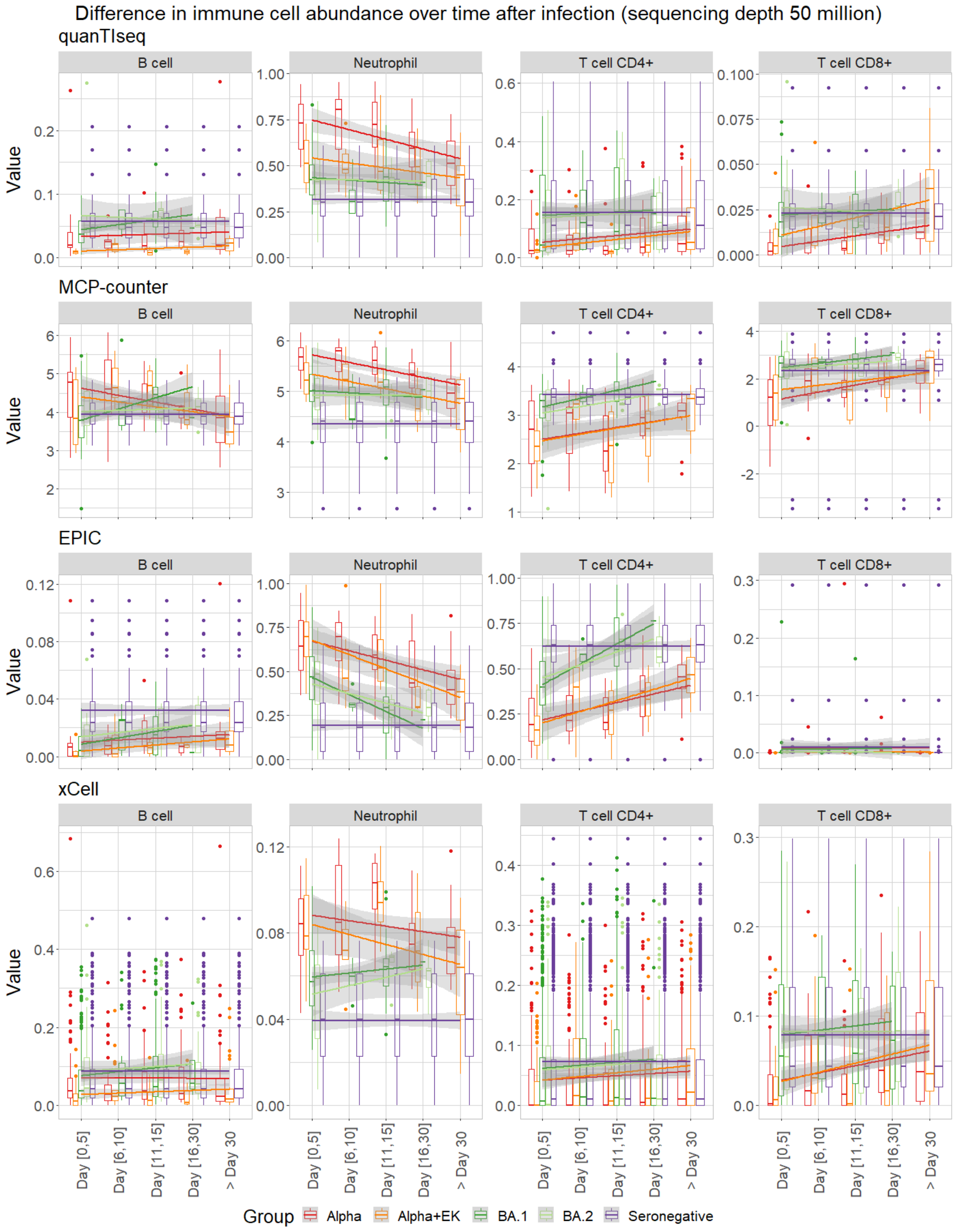


**Supplemental Figure 6a:** Immune deconvolution results for downsampled sequencing depth 10 million. Cell-type fractions separated over brackets 0-5, 6-10, 11-15, 16-30, and >30 days after hospitalization or onset of symptoms detected by the immune deconvolution methods quanTseq, MCP-counter, EPIC, and xCell for the immune cells B cell, Neutrophil, T cell CD4+, and T cell CD8+. The Gamma variant has been removed in this analysis due to poor sample size per time bracket (Suppl. Tables 1a-b).

### Supplemental Figure 6b:

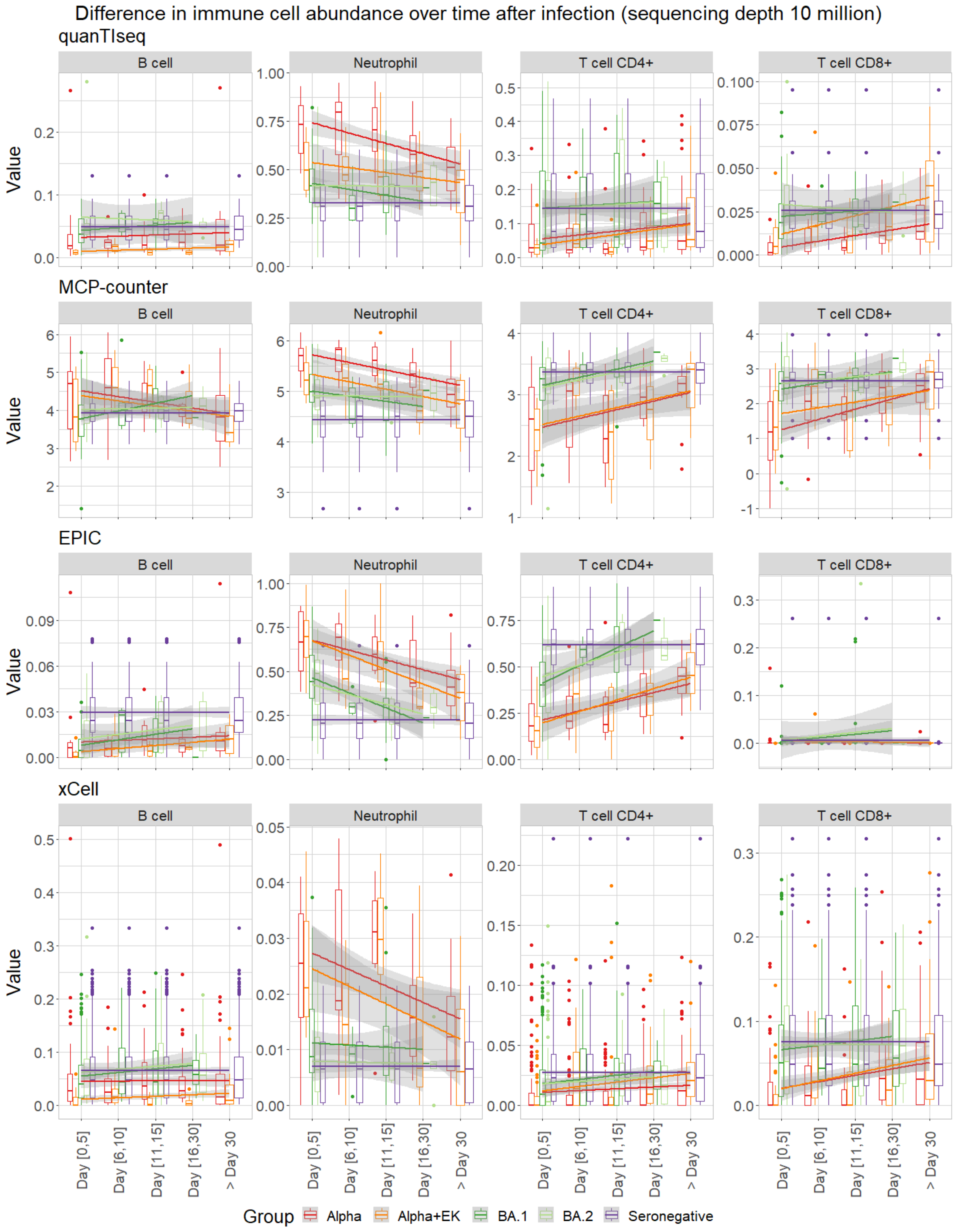


**Supplemental Figure 6b:** Immune deconvolution results for downsampled sequencing depth 10 million. Cell-type fractions separated over brackets 0-5, 6-10, 11-15, 16-30, and >30 days after hospitalization or onset of symptoms detected by the immune deconvolution methods quanTseq, MCP-counter, EPIC, and xCell for the immune cells B cell, Neutrophil, T cell CD4+, and T cell CD8+. The Gamma variant has been removed in this analysis due to poor sample size per time bracket (Suppl. Tables 1a-b).

**Supplemental Figure 7a: Correlation of experimental complete blood count of Neutrophils, Lymphocytes (B cells, T cells), and Monocytes for sequencing depth 50 Million.**

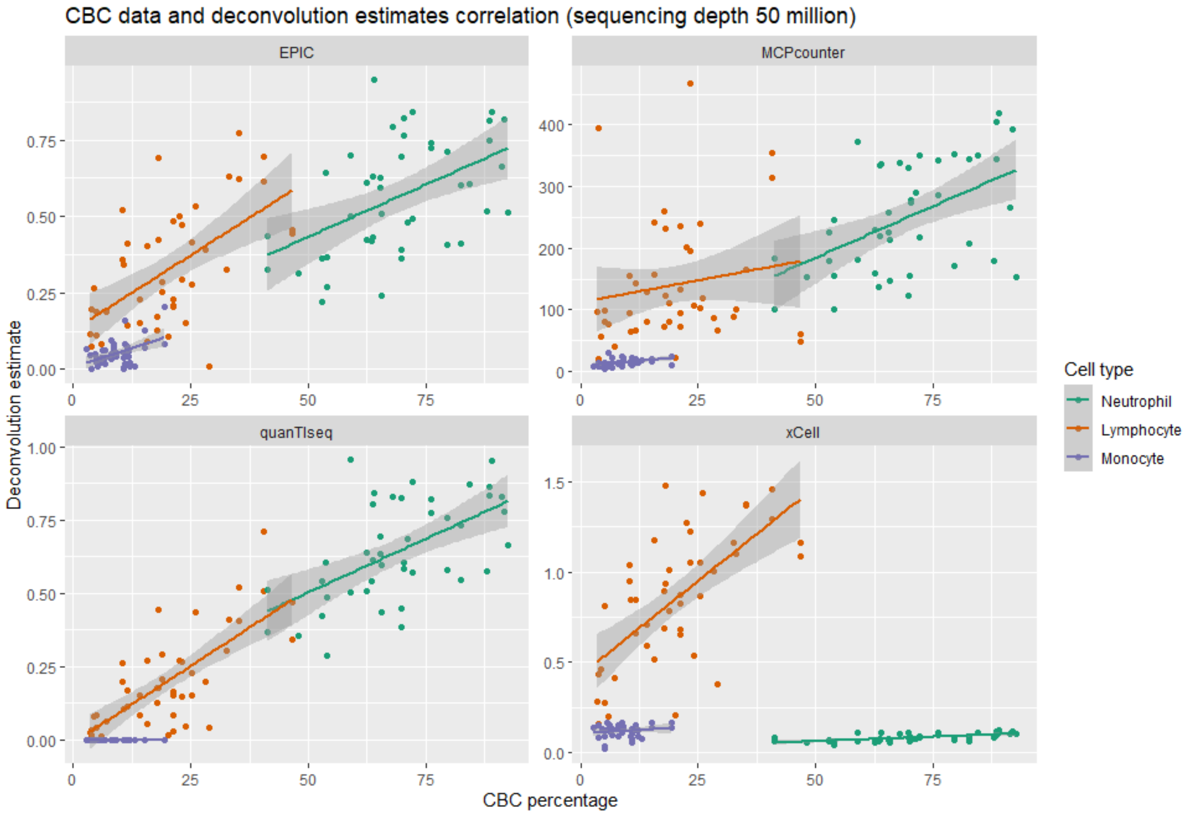


**Supplemental Figure 7a**: Correlation between CBC data and immune deconvolution scores for a sequencing depth of 10 million across Lymphocytes, Monocytes, and Neutrophils using all four deconvolution methods. *Statistical values: EPIC: Neutrophil: R=0.49, p=0.00082, RMSE=70.07, Lymphocyte: R=0.58, p=3.2e-05, RMSE=22.74, Monocyte: R=0.48, p=0.001, RMSE=9.71; MCPcounter: Neutrophil: R=0.5, p=0.00058, RMSE=198.32, Lymphocyte: R=0.17, p=0.28, RMSE=154.05, Monocyte: R=0.43, p=0.0036, RMSE=7.55; quanTIseq: Neutrophil: R=0.57, p=6.2e-05, RMSE=69.99, Lymphocyte: R=0.73, p=1.4e-08, RMSE=22.84, Monocyte: R=NA, p=NA, RMSE=9.77; xCell: Neutrophil: R=0.62, p=8.3e-06, RMSE=70.57, Lymphocyte: R=0.65, p=1.4e-06, RMSE=22.21, Monocyte: R=0.17, p=0.27, RMSE=9.66.*

**Supplemental Figure 7b: Correlation of experimental complete blood count of Neutrophils, Lymphocytes (B cells, T cells), and Monocytes for sequencing depth 10 Million.**

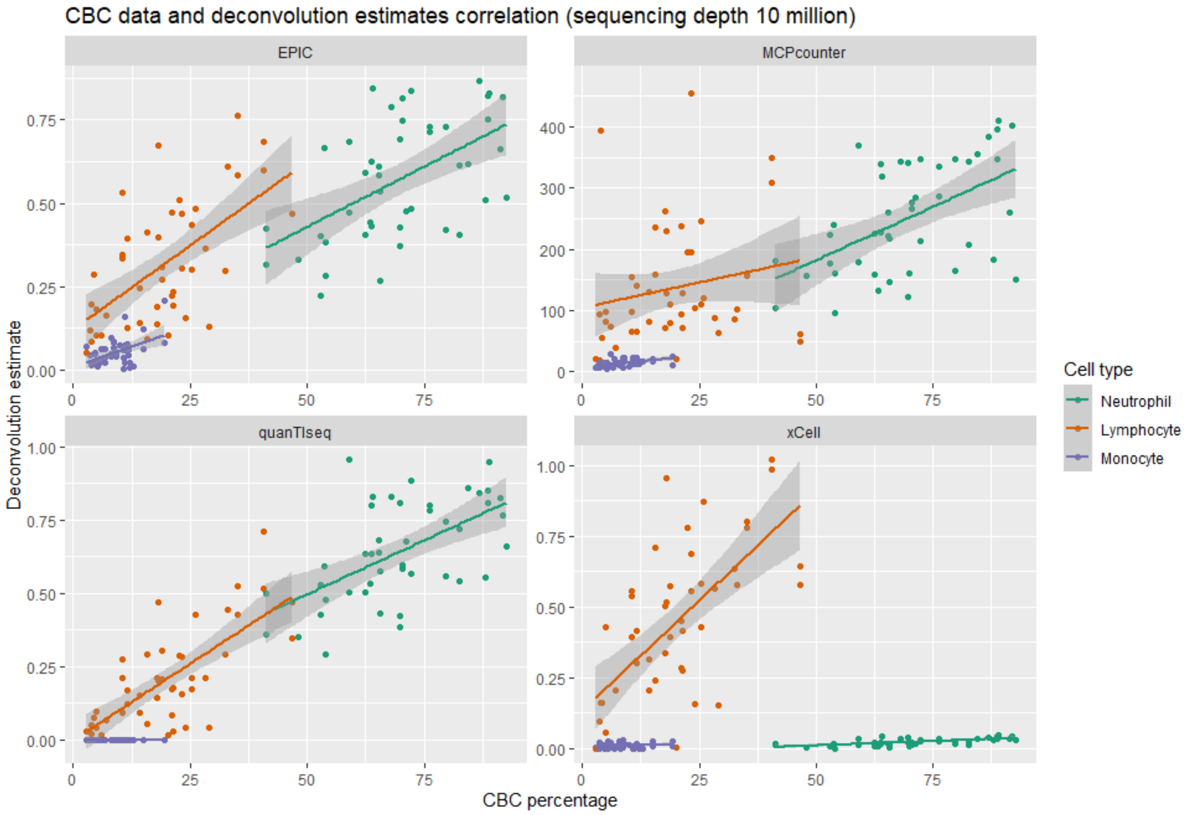


**Supplemental Figure 7b**: Correlation between CBC data and immune deconvolution scores for a sequencing depth of 10 million across Lymphocytes, Monocytes, and Neutrophils using all four deconvolution methods. *Statistical values: EPIC: Neutrophil: R=0.54, p=0.00016, RMSE=70.48, Lymphocyte: R=0.63, p=3.4e-06, RMSE=22.49, Monocyte: R=0.47, p=0.0012, RMSE=9.71; MCPcounter: Neutrophil: R=0.52, p=0.00025, RMSE=200.05, Lymphocyte: R=0.2, p=0.19, RMSE=151.05, Monocyte: R=0.45, p=0.0018, RMSE=7.76; quanTIseq: Neutrophil: R=0.58, p=3.9e-05, RMSE=70.41, Lymphocyte: R=0.73, p=9.5e-09, RMSE=22.59, Monocyte: R=NA, p=NA, RMSE=9.77; xCell: Neutrophil: R=0.66, p=9.5e-07, RMSE=71.03, Lymphocyte: R=0.66, p=7.4e-07, RMSE=22.35, Monocyte: R=0.24, p=0.11, RMSE=9.76.*

**Supplemental Figure 8:**

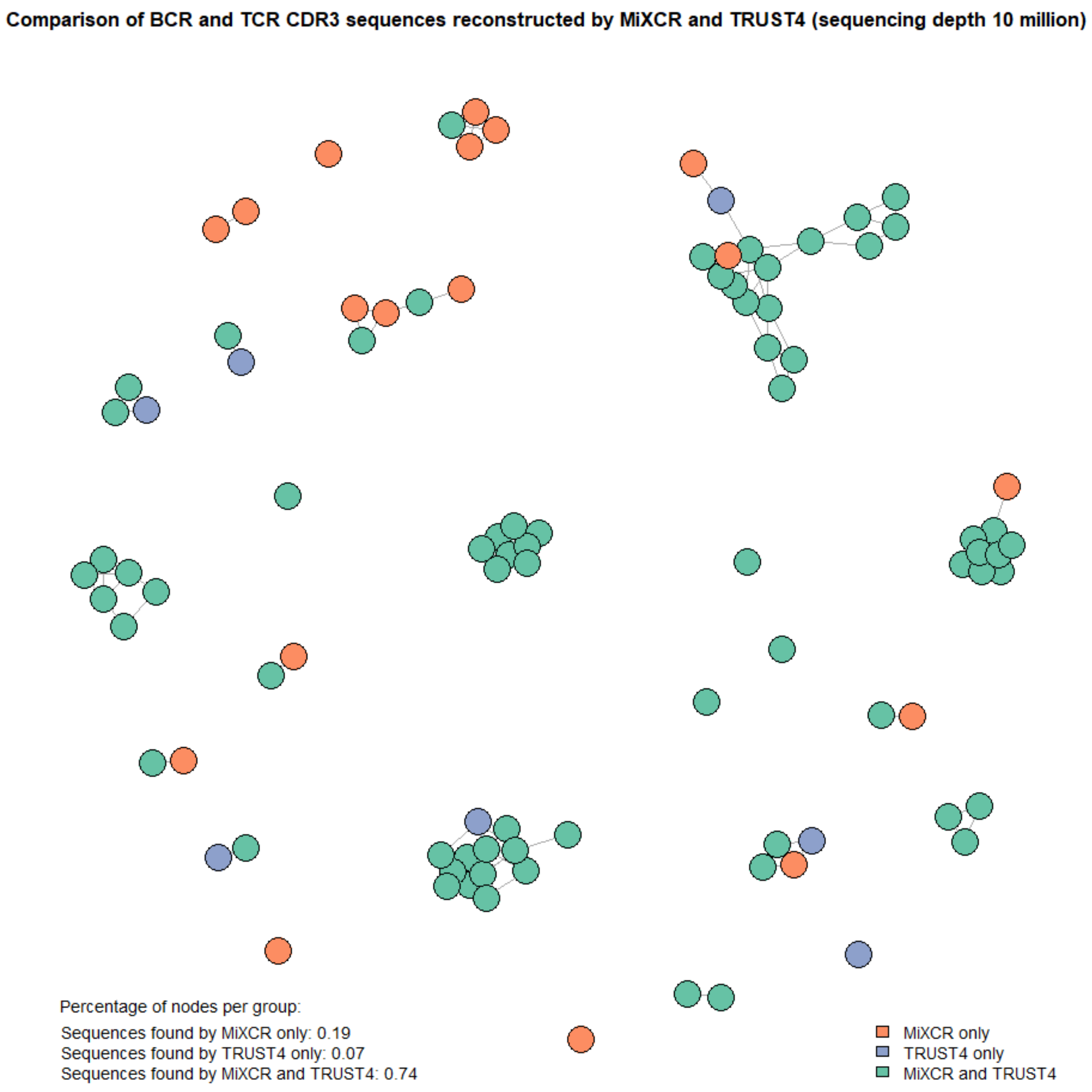


**Supplemental Figure 8:** Network displaying the BCR and TCR sequences reconstructed by MiXCR (orange) and TRUST (blue) and sequences found by both tools as nodes. Interactions (edges) between the sequences show sequence similarities. 74 % of unique sequences were found by both tools, 19 % were only found by MiXCR, and 7 % were only found by TRUST4.

### Supplemental Figure 9:

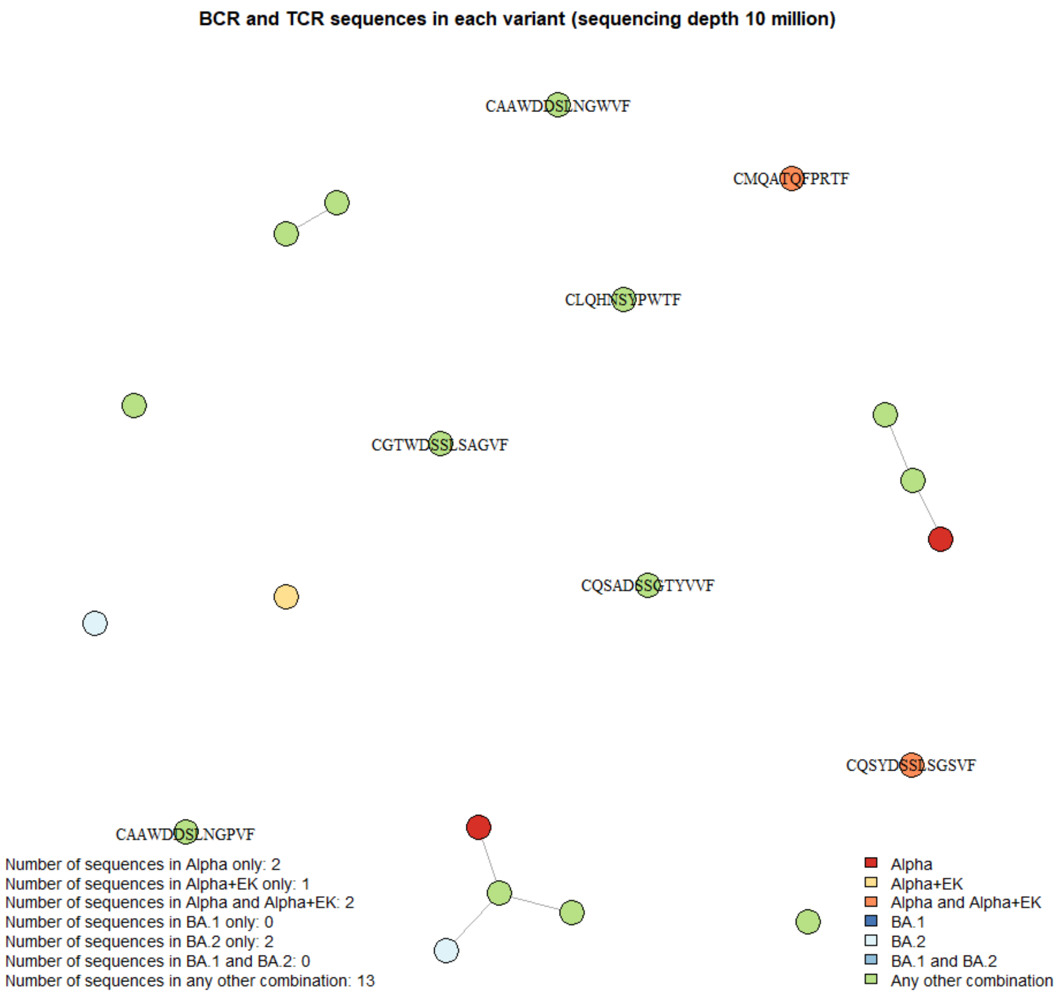


**Supplemental Figure 9:** Network showing sequences that are only found in infected samples as nodes and sequence similarities (score below 10) as edges. Annotated sequences are sequences with an anti-SARS-CoV-2 immunoglobulin pBLAST hit.
